# Exocyst subunit Sec6 is positioned by microtubule overlaps in the moss phragmoplast prior to the arrival of cell plate membrane

**DOI:** 10.1101/368860

**Authors:** Han Tang, Jeroen de Keijzer, Elysa Overdijk, Els Sweep, Maikel Steentjes, Joop E. Vermeer, Marcel E. Janson, Tijs Ketelaar

## Abstract

During plant cytokinesis a radially expanding membrane-enclosed *cell plate* is formed from fusing vesicles that compartmentalizes the cell in two. How fusion is spatially restricted to the site of cell plate formation is unknown. Aggregation of cell-plate membrane starts near regions of microtubule overlap within the bipolar phragmoplast apparatus of the moss *Physcomitrella patens*. Since vesicle fusion generally requires coordination of vesicle tethering and subsequent fusion activity we analysed the subcellular localization of several subunits of the exocyst, a tethering complex active during plant cytokinesis. We found that Sec6, but neither Sec3 or Sec5 subunits localized to microtubule overlap regions in advance of cell plate construction started in moss. Moreover, Sec6 exhibited a conserved physical interaction with an orthologue of the Sec1/Munc18 protein KEULE, an important regulator for cell-plate membrane vesicle fusion in Arabidopsis. Recruitment of PpKEULE and vesicles to the early cell plate was delayed upon Sec6 gene silencing. Our findings thus suggest that vesicle-vesicle fusion is in part enabled by a pool of exocyst subunits at microtubule overlaps that is recruited independent of the delivery of vesicles.

**Summary statement:** We performed a time-resolved localization screen of multiple subunits of the exocyst complex throughout moss cytokinesis and show that each subunit has a unique spatiotemporal recruitment pattern.

## Introduction

The physical separation of the two daughter cells formed during plant cell division occurs via a transient, disk-shaped membrane compartment that expands radially towards the parental cell wall. This membrane compartment is termed the cell plate and its construction culminates in a new cell wall segment dividing two individual plasma membranes (Drakakaki, 2015; Müller and Jürgens, 2015; Smertenko et al., 2017). The initiation of the cell plate and its radial expansion relies on the fusion of vesicles that are supplied mainly by the secretory pathway (Mcmichael and Bednarek, 2013; Richter et al., 2014; Boruc and Van Damme, 2015). Adaptations of canonical trafficking mechanisms are however required because there is no pre-existing target membrane at the site of cell division to which vesicles can fuse. Instead, membrane deposition is thought to be initiated by ‘homotypic’ fusion of vesicles (Smertenko et al., 2017). This raises the question of how vesicle fusion is spatially restricted to the site of cell division and not spuriously throughout the cell when vesicles meet. It is thought that vesicles are transported along polarized microtubules to the centre of the phragmoplast, a cytoskeletal apparatus that supports cell plate assembly. Although locally concentrating vesicles may enhance the fusion rates of fusion-competent vesicles, it is unclear whether a transport-based mechanism alone can provide the spatial accuracy required to build a straight and flat cell plate. Recently we identified short stretches of antiparallel microtubule overlap at the midzone of phragmoplasts in the moss *Physcomitrella patens* as sites where membrane build up is initiated (de Keijzer et al., 2017). It remained however unclear whether there are molecules at overlaps that trigger vesicle fusion locally.

In eukaryotic cells the fusion of transport vesicles with endomembrane compartments and the plasma membrane relies on the combined action of fusion and tethering complexes. The force driving the fusion of a vesicle and its destination membrane is almost universally generated by SNARE (soluble *N*-ethylmaleimide-sensitive factor attachment receptor) complexes, typically composed of 4 conserved, membrane-associated proteins (Söllner et al., 1993; Wickner and Schekman, 2008; Südhof and Rothman, 2009). Tethering – the establishment of the initial physical connection between the two membranes – on the other hand, is orchestrated by a multitude of molecular machinery, often in form of multimeric protein complexes (Koumandou et al., 2007; Vukašinović and Žárský, 2016). To enable targeted trafficking among the various distinct membrane compartments in eukaryotic cells, membranes acquire different identities that dictate which membrane fusion reactions are allowed. Although SNARE complex composition bestows some of this specificity (Paumet et al., 2004), other factors including Rab GTPases (Grosshans et al., 2006; Stenmark, 2009) and the various tethering complexes play an important role as well (Yu and Hughson, 2010). Indeed, each of the several tethering complexes typically facilitates the docking of vesicles at a specific target membrane (Vukašinović and Žárský, 2016; Yu and Hughson, 2010). While work in Arabidopsis has uncovered much about the composition, trafficking and function of SNARE complexes involved in fusing cell plate vesicles (reviewed in Müller and Jürgens, 2015; Jürgens et al., 2015), comparatively little is known about the tethering factors involved. In Arabidopsis both the TRAPPII (Transport Protein Particle II) and exocyst tethering complexes contribute to cell plate biogenesis (Thellmann et al., 2010; Fendrych et al., 2010; Qi et al., 2011; Rybak et al., 2014). Both are multimeric protein complexes that function in post-Golgi membrane trafficking (Drakakaki et al., 2012; Fendrych et al., 2010; reviewed by Vukašinović and Žárský, 2016). Interestingly, while TRAPPII is associated with the cell plate throughout its formation, the exocyst is present on the cell plate during early membrane fusion, whereafter its abundance drops during cell plate extension and increases again after cell plate insertion (Fendrych et al., 2010; Zhang et al., 2013; Rybak et al., 2014). Nonetheless, the two tethering complexes physically interact and most likely function cooperatively during cytokinesis (Rybak et al., 2014). While functional analysis of the TRAPPII complex is expedited by only single genes encoding the two unique TRAPPII subunits (Thellmann et al., 2010; Qi et al., 2011), the diverse complement of genes encoding exocyst subunits have made dissecting the role of the exocyst comparatively cumbersome. Nonetheless, various exocyst mutants have been identified that harbour cell plate defects (Fendrych et al., 2010; Wu et al., 2013; Rawat et al., 2017).

Here, to understand if the exocyst could be a cell-plate assembly factor active on microtubule overlaps, we monitored the localization of several GFP-tagged exocyst subunits during cytokinesis in the moss *P. patens*. Although this representative of a basal land plant lineage has seen a similar degree of gene amplification of exocyst subunit genes as Arabidopsis, with a notable smaller radiation of Exo70 paralogs (Cvrčková et al., 2012), it features a major haploid phase in its life cycle expediting genetic analysis. Since cell plate assisted cytokinesis is a hallmark of all land plants (Buschmann and Zachgo, 2016) and a comprehensive toolset to study cell division exists for *P. patens* (Yamada et al., 2016), we anticipated that this model plant is particularly useful for studying exocyst function in relation to cell plate membrane fusion. We show that exocyst subunit Sec6, and not the subunits Sec3 and Sec5, localizes to microtubule overlaps at the phragmoplast midzone prior to the arrival of membrane vesicles. The localisation of Sec6 is dependent on the presence and size of the microtubule overlaps. Our findings make the case that, in general, spatial control over homotypic vesicle fusion can be achieved by linking tethering activity to a pre-defined, non-membranous subcellular structure, which during plant cytokinesis are microtubule overlaps.

## Results

In eukaryotes, the exocyst facilitates targeting and fusion of exocytotic vesicles to the plasma membrane. A key step in exocyst functioning is the formation of a fully assembled complex when contact is made between a subset of vesicle-associated exocyst subunits and plasma-membrane associated subunits (e.g. Yu and Hughson, 2010; Boyd et al., 2004). How exocyst assembly proceeds in absence of a plasma membrane in the context of homotypic fusion is unknown. To characterize assembly dynamics during cytokinesis in *P. patens*, we generated a time resolved localization map of selected exocyst subunits in protonemal tip cells that exhibit repetitive cell divisions. Sec6 was included in the screen because it is the only exocyst subunit that is represented by a single gene in the moss *P. patens* (Cvrčková et al., 2012). The protein is therefore expected to be part of all fully assembled exocyst complexes. Moreover Sec6 from Arabidopsis was shown to interact with vesicle fusion machinery and may thus be a regulator of cell plate formation (Wu et al., 2013). To investigate whether exocyst complex assembly occurs in sequential steps we included Sec5 and Sec3 paralogs, which in yeast were proposed to be vesicle and plasma membrane associated respectively (He and Guo, 2009). Sec5 is encoded by 4 genes and Sec3 by 3 genes in *P. patens*. To conduct the screen, a GFP-encoding fragment was integrated at the end of the single Sec6 gene, all three genes encoding Sec3 paralogs, and the three Sec5 paralogs that are expressed in the protonemal tissue (Ortiz-Ramírez et al., 2016) (Figure S1A). Firstly we analysed the relative abundance of the different isoforms by imaging interphase apical caulonemal cells (Figure S1B). The microscopical assessment of relative protein abundance followed the gene expression data in S1A. All investigated exocyst subunits with detectable amounts of fluorescence localized to plasma membrane foci as reported previously (Fendrych et al., 2013; Zhang et al., 2013; Vukašinović and Žárský, 2016; Bloch et al., 2016; Synek et al., 2017; an Gisbergen et al., 2018). Of thesubunits that showed expression levels suitably high for microscopy, fusion proteins were expressed alongside mCherry-α-tubulin such that exocyst localization dynamics could be related to mitotic progression (Figure S1, 1A). All selected exocyst subunits and additionally an established cell plate membrane marker (SCAMP4; de Keijzer et al. 2017) were imaged throughout cell division. The onset of anaphase was used as temporal reference to aid in the mutual comparison of arrival times to specific cytokinetic structures (Figure 1A, Movie S1).

**Figure 1.**
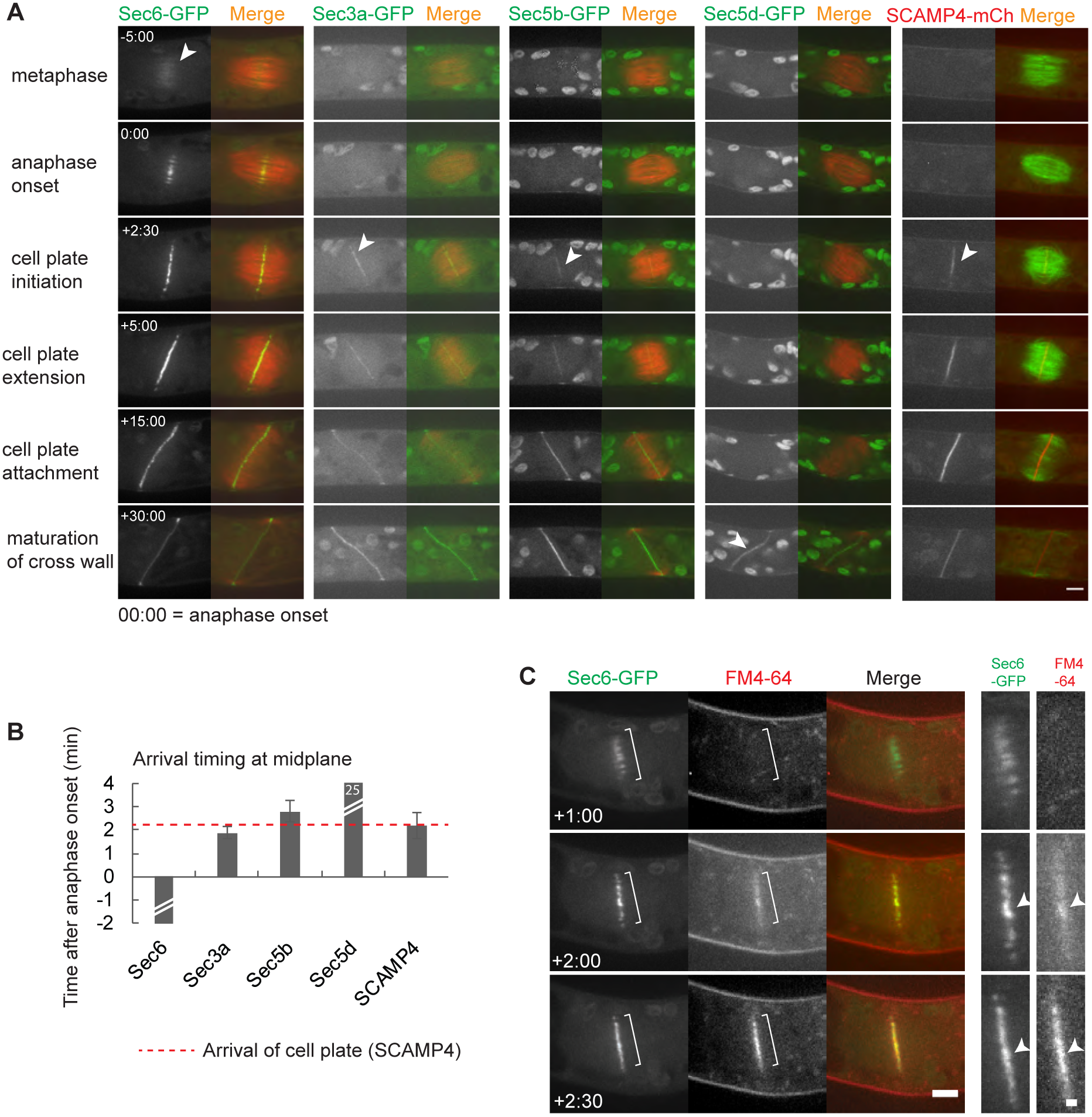
Localization survey of a subset of *Physcomitrella patens* exocyst subunits during cell division. (A) Localization of exocyst subunits Sec6, Sec3a, Sec5b and Sec5d during cell division visualized in caulonemal apical cells expressing mCherry-α-tubulin (appearing red in the merged image) and a GFP-tagged version of the indicated exocyst subunit. The progression of cell plate and phragmoplast development, visualized with cell plate marker SCAMP4 (de Keijzer et al. 2017) and GFP-α-tubulin, is depicted on the right as a reference. Time after anaphase onset (t=0) is indicated in min:sec. Localization to the phragmoplast midzone is indicated with white arrowheads. Images are maximum z-projections of 3 confocal planes spaced 0.5 μm apart. Scalebar, 5 μm. (B) Bar graph showing the average appearance time of GFP-tagged exocyst subunits at the phragmoplast midzone compared to that of cell plate membrane (SCAMP4-mCherry, dashed line). Error bars indicate standard deviation. Averages were obtained from n=4 cells (Sec6 and Sec3a), n=5 cells (Sec5b and Sec5d) or n=8 cells (SCAMP4). (C) Early cell plate membrane accumulation visualized in a dividing cell expressing Sec6-GFP stained with FM4-64 membrane dye. Time with respect to anaphase onset (t=0) is indicated in min:sec. The bracketed areas are shown in detail on the right. Here, arrowheads mark membranous material accumulating at Sec6-labelled sites. Images are maximum z-projections of 3 confocal planes spaced 0.5 μm apart. Scalebar in overview images, 5 μm; Scalebar in zoomed images, 1 μm. Sec6 resides to microtubule overlaps at mitotic onset prior to other subunits that arrive later conccurent with vesicles gathering during cytokinesis.

The selected exocyst subunits showed disparate localization patterns during cell division. Sec3a and Sec5b localized to the phragmoplast midzone during cell plate initiation (Figure 1A, Movie S1). Unlike the exocyst subunits studied in Arabidopsis (Fendrych et al., 2010; Zhang et al., 2013; Rybak et al., 2014), no sharp drop in intensity occurred during cell plate expansion. After cell plate attachment to the parental wall, similar to the observations in Arabidopsis, the amount of Sec3a and Sec5b on the new wall facet transiently increased. In contrast, Sec5d deviated from this localization and was absent from the cell plate until late cytokinesis. Moreover, Sec6 was detected already at the centre of the division apparatus from spindle formation onwards (Figure 1A, white arrowheads; Movie S1), long before the arrival of cell plate membrane (Figure 1C, Movie S2). Unlike the other studied exocyst subunits, the intensity of Sec6 peaked during the onset of cytokinesis and cell plate expansion, where after it dropped after cell plate attachment to the parental wall. Interestingly, all studied subunits associated with the cell cortex already before cell plate attachment at a site where the division plane intersected with the parental cell wall (Figure 2A and 2B). To our knowledge such a cortical localization pattern has not been observed during cytokinesis for any labelled subunit in Arabidopsis (Fendrych et al., 2010; Zhang et al., 2013; Rybak et al., 2014). The early cortical labelling was punctate, with numerous puncta together making up a 2-3 μm wide band that formed a continuous ring at the cell cortex (Figure 2A). The band was mobile on the cortex until cell plate attachment and its movement was synchronous with the rotation of the phragmoplast in these cells (Figure 2B and 2D.

**Figure 2.**
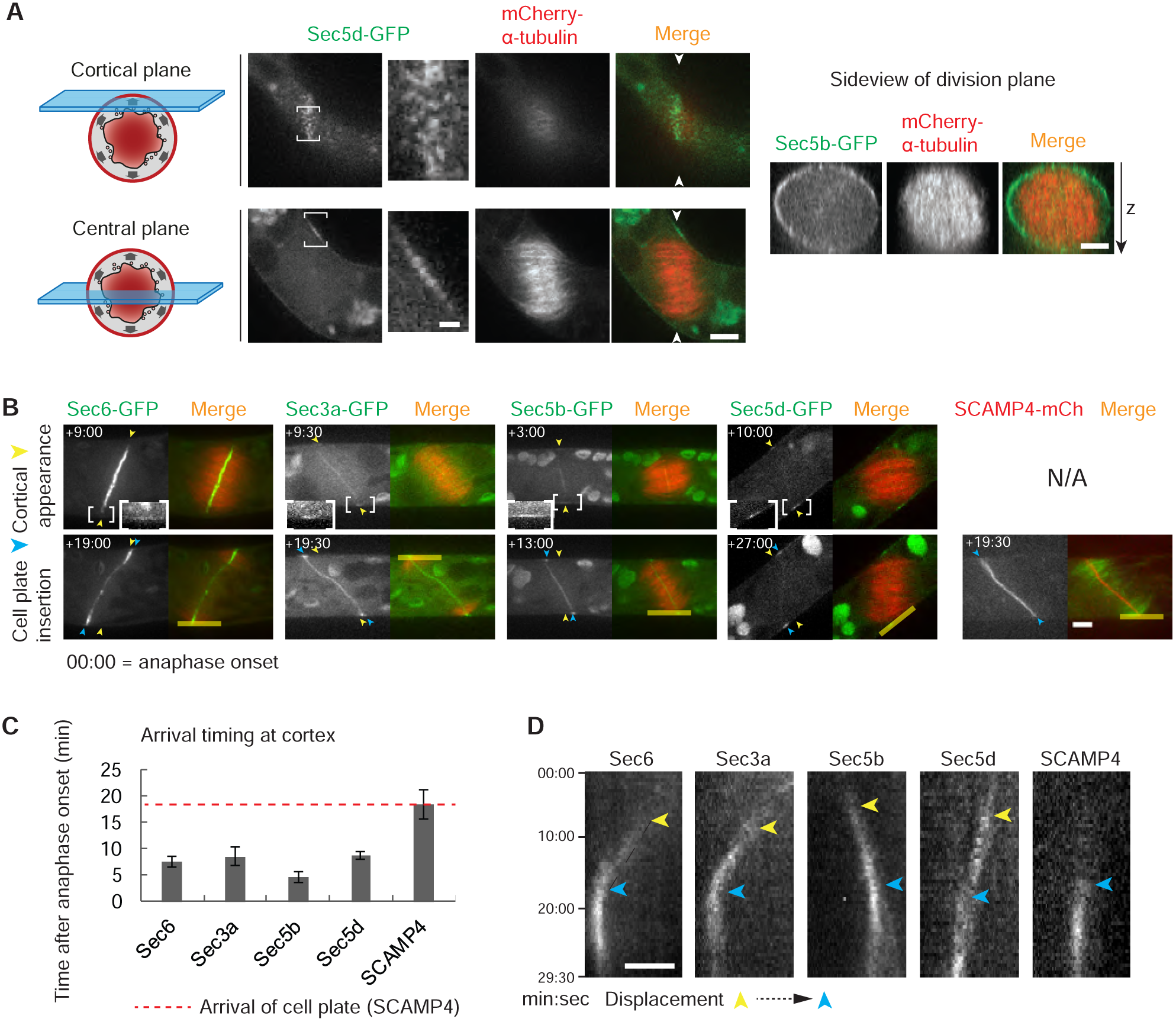
Exocyst subunits arrive to cortical membrane prior to cell plate insertion. (A) Cortical localization of the exocyst during cell division visualized in a cell expressing Sec5d-GFP and mCherry-α-tubulin at a single cortical and central confocal plane (illustrated on the left). A detailed view of the Sec5d-GFP signal indicated by the brackets is shown. A sideview of the division plane (marked with arrowheads) is depicted on the right. Scalebar in overview and sideview images, 5 μm; Scalebar in zoomed images, 1 μm. (B) Snapshots of dividing caulonemal cells expressing exocyst subunit-GFP (Sec6, Sec3a, Sec5b and Sec5d), and membrane marker SCAMP4-mCherry. Microtubules labelled with mCherry-α-tubulin or GFP were used as a temporal reference. Images are maximum z-projections of 3 planes spaced 0.5 μm apart acquired in the central plane. Images were recorded with 30 sec interval. Time with respect to anaphase onset (t=0) is indicated in min: sec. Scale bar=5 μm. The time point each subunit first appeared at the cortex is indicated by yellow arrowheads in the upper panels, which have been copied to the lower panels. The bracketed areas are shown in detail. The final localization of each subunit in the cortex at the moment of cell plate insertion is indicated by blue arrowheads in the lower panel. (C) Bar graph showing the average appearance time of GFP-tagged exocyst subunits at the cell cortex compared to that of cell plate membrane (SCAMP4-mCherry, dashed line). Error bars indicate standard deviation. Averages were obtained from n=4 cells (Sec6 and Sec3a), n=5 cells (Sec5b and Sec5d) or n=8 cells (SCAMP4). (D) Kymograph of exocyst subunits and SCAMP4 generated along the parental plasma membrane as indicated by the yellow lines depicted in B. A region of 0.67 μm perpendicular to the line was used to calculate average intensity. The yellow arrowheads mark the timing of arrival while the blue arrowheads indicate the timing of cell plate insertion.

To further characterize the observed differences in localization, we identified the moment with respect to anaphase onset at which subunits first became visible at both the phragmoplast midzone and the cortex (Figure 1B and 2C). This analysis showed that all studied subunits arrived at the cortex at around the same time (5-10 minutes post anaphase onset). The average time of arrival at the midzone however diverged enormously for the studied subunits. Since the dynamics of exocyst subunits recruitment during cytokinesis depends on the cellular location, regulatory mechanisms for exocyst recruitment to the phragmoplast midzone and the cortex must be in place. To exclude that the weakly expressing Sec3b-GFP and Sec5a-GFP behaved differently during cytokinesis than the other Sec3 and Sec5 paralogs, we imaged cytokinesis in these lines over time after application of the membrane marker FM4-64 (De Keijzer et al., 2017).

This showed that these weakly expressing paralogs had similar recruitment dynamics as the other Sec3 and Sec5 paralogs (Figure S1C). Thus, while sharing localization at the cortex and late cell plate with the other studied exocyst subunits, Sec6 also exhibited localization to the phragmoplast and spindle midzone apparently independent of the other studied exocyst complexes and membrane compartments.

The early localization pattern of Sec6 showed strong resemblance to the sites where microtubules from opposite poles form antiparallel overlaps in the spindle and phragmoplast midzone (Ho et al., 2011; Kosetsu et al., 2013; de Keijzer et al., 2017). To investigate their interdependency we generated a moss line expressing Sec6-mCherry together with a Citrine-tagged version of the antiparallel microtubule bundling protein MAP65 (Kosetsu et al., 2013). Sec6 localization indeed closely followed the distribution of MAP65-citrine labelled regions of overlap (Figure 3, left panels; Movie S3). Strikingly, this behaviour was even visible during prophase, when the nuclear envelope is still intact and no bipolar spindle is yet formed (Figure 3, arrowheads). We previously generated cells lacking a regulator of microtubule dynamics, Kin4-Ic, that show impaired overlap shortening at anaphase onset (de Keijzer et al., 2017). In this genetic background a similar strong co-localization of MAP65 and Sec6 was observed during prophase and spindle stages, yet Sec6 increasingly localized to only the central region of elongated overlaps as cytokinesis progressed (Figure 3, right panels; Movie S3). Nevertheless, during early cytokinesis the width of the Sec6 patches was wider in Δ*kin4-Ic* cells compared to wild-type (Figure 3, arrows). The localization of Sec6 along overlaps showed high resemblance with membrane depositions which also gradually became more confined to overlaps centers (de Keijzer et al., 2017). Thus, Sec6 localizes to antiparallel overlaps but during cell plate construction other factors may additionally affect its localization.

**Figure 3.**
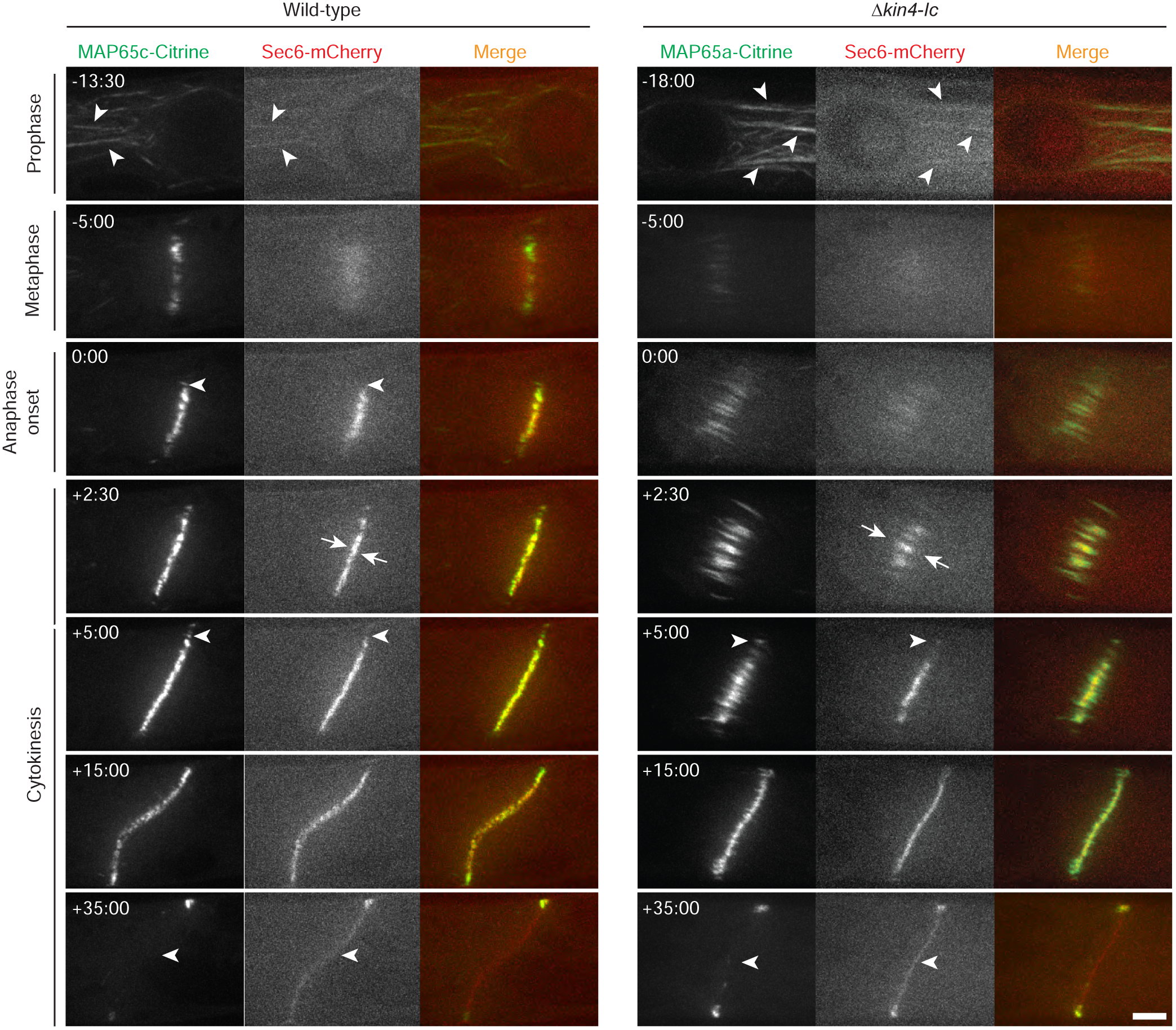
Exocyst subunit Sec6 partially co-localizes with MAP65 on antiparallel microtubule overlaps. Caulonemal cells expressing citrine labelled MAP65 and Sec6-mCherry in a wild-type and Δ*kin4-Ic* genetic background imaged throughout mitosis. MAP65-labelled microtubule bundles present during prophase that show concomitant Sec6 labelling are marked with arrowheads. Arrows highlight the width of the Sec6-labelled zone during early cytokinesis. Time with respect to anaphase onset (t=0) is indicated in min:sec. Images are maximum z-projections of 3 confocal planes spaced 0.5 μm apart. Scalebar, 5 μm.

To further understand to what extent the presence of microtubule overlaps are a prerequisite for the localization of Sec6 during cytokinesis, we silenced the three MAP65 genes expressed in protonema. Earlier studies have demonstrated that upon knockdown of MAP65 antiparallel microtubule overlap formation in the phragmoplast is severely compromised, ultimately leading to phragmoplast collapse. Nonetheless, during early cytokinesis cell plate membrane aggregates and phragmoplast microtubules remain organized as two bipolar sets, but with compromised overlap formation in the centre (Kosetsu et al., 2013). While in control cells the Sec6-GFP appeared regularly distributed over the division plane during early cytokinesis, in MAP65 silenced cells Sec6-GFP formed discontinuous patterns (Figure 4; Movie S4). There was no obvious correlation between the appearance of gaps in the Sec6 localization pattern and the presence of microtubules; areas devoid of Sec6 were visible both in regions populated and unpopulated by microtubules (Figure 4). Localization of Sec6 to the phragmoplast midzone therefore not just correlates with the presence of bipolar microtubules but requires an ordered array of microtubule overlaps.

**Figure 4.**
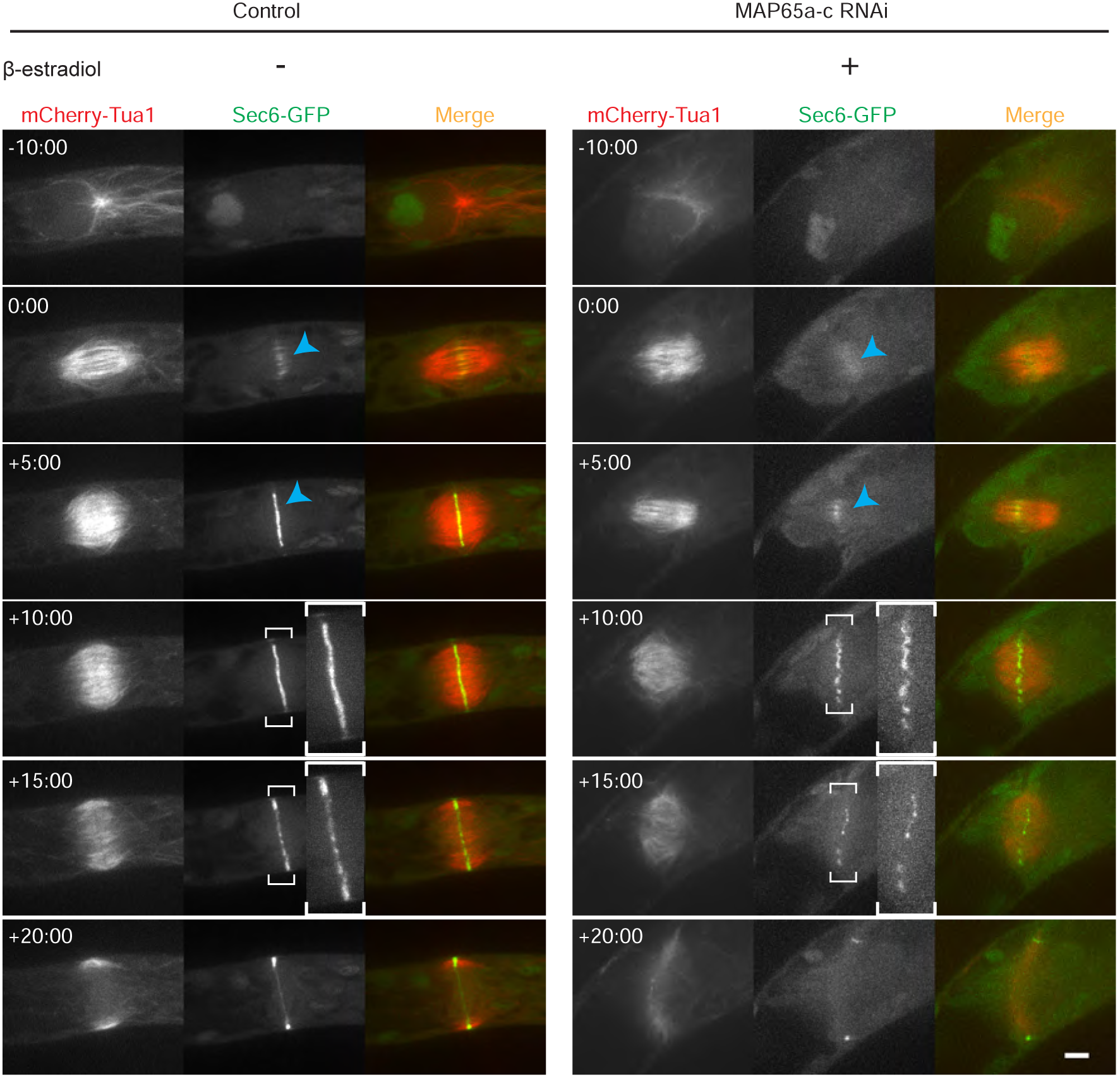
The localization of exocyst subunit Sec6 to antiparallel microtubule overlaps depends on overlap length and presence of MAP65. Time sequence of dividing caulonemal cells expressing Sec6-GFP and mCherry-α-tubulin with silencing of MAP65a, b and c non-induced (control) and induced though the addition of β-estradiol. Arrowheads mark the accumulation of Sec6 to the phragmoplast midzone during early cytokinesis and the bracketed areas delineate the midzone during a later stage of cytokinesis in detail. Time with respect to anaphase onset (t=0) is indicated in min:sec. Images are maximum z-projections of 3 confocal planes spaced 0.5 μm apart. Scalebar, 5 μm. MAP65 is required for SEC6 accumulation to microtubule overlaps.

A physical interaction between Sec6 and Sec1/munc18 (SM) family protein KEULE was proposed to be an important regulatory step in cytokinetic vesicle fusion in Arabidopsis (Wu et al., 2013). KEULE contributes to fusion by preventing the important cytokinetic SNARE-component KNOLLE from refolding into its closed, non-fusion-competent conformation, thereby allowing the formation of fusogenic trans-SNARE complexes among vesicles delivered to the cell plate (Park et al., 2012; Karnahl et al., 2017; Jürgens et al., 2015). Whether the Sec6 interaction regulates this specific activity is presently unknown. Nonetheless, in Sec6-mutants cytokinesis defects were found that resemble these encountered in plants lacking KEULE (Wu et al., 2013; Assaad et al., 1996), hinting that a cooperative Sec6-KEULE interaction regulates cytokinesis. Based on these findings we hypothesized that Sec6 on overlaps may regulate vesicle fusion activity in space through an interaction with KEULE. In search of a KEULE homologue we identified 7 loci in the *P. patens* genome that are predicted to encode SM family proteins. Of the identified loci, two contained genes with a predicted exon number equal to that of the Arabidopsis KEULE gene and their expected gene products exhibited an overall similarity to Arabidopsis KEULE at least twice higher than the other predicted SM genes (Figure 5A). Analysis of gene expression levels revealed that the gene encoded at locus Pp3c17_24130 was ubiquitously expressed in all moss tissues, while the other gene (Pp3c16_570) was only expressed in rhizoids (Figure 3A and Figure S2; Ortiz-Ramírez et al., 2016). We therefore focussed on the former gene and tentatively named it PpKEULE. Since the position of introns in genes contains information on the evolutionary trajectory of the gene family they belong to (e.g. Rogozin et al., 2003; Garcia-España et al., 2009; Javelle et al., 2011), we compared the intron-exon structure between Arabidopsis KEULE and PpKEULE. The intron positions were highly similar for both species (Figure 5B).

**Figure 5.**
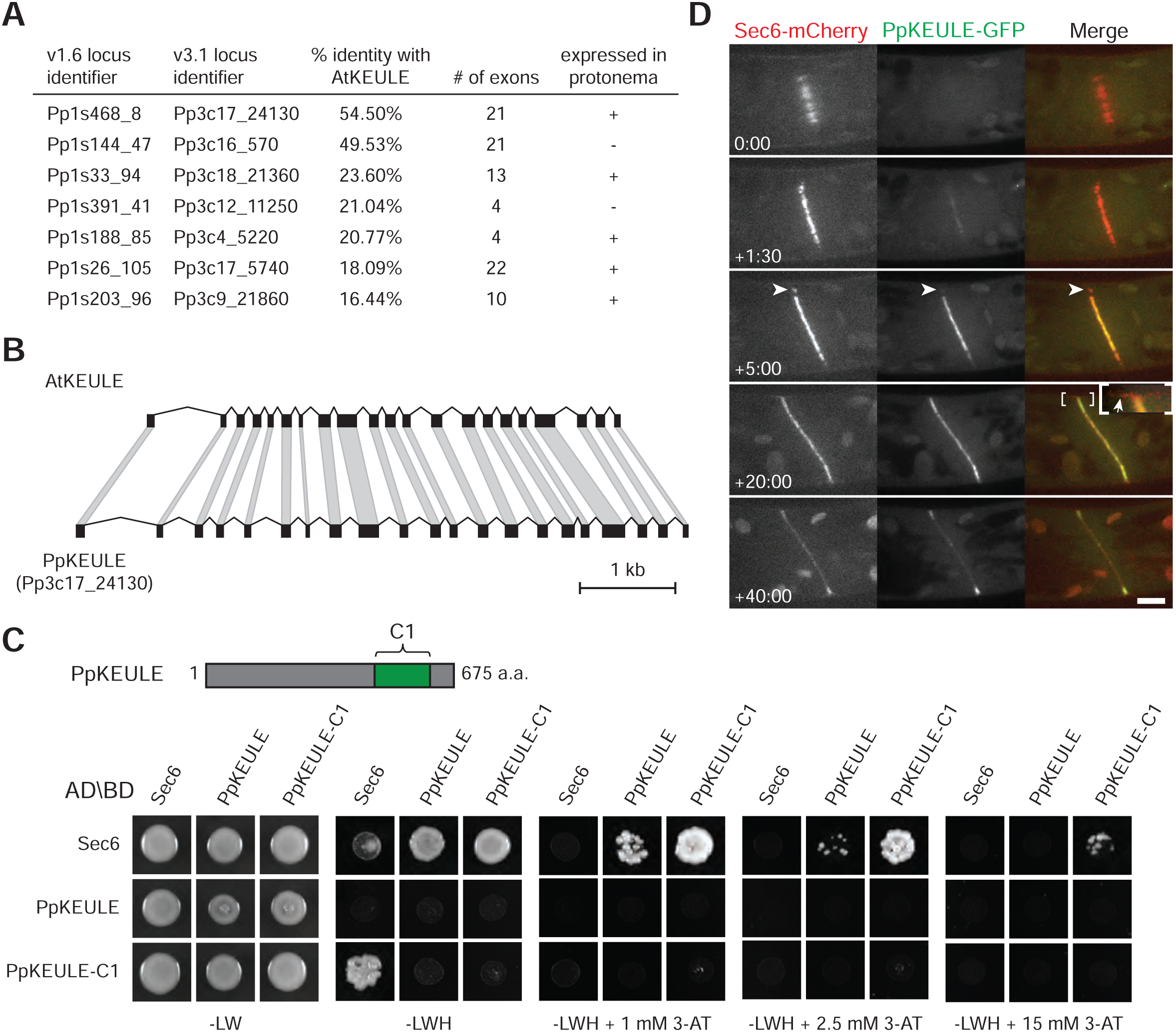
Identification of a *P. patens* KEULE orthologue that physically interacts with Sec6 and co-localizes at the phragmoplast midzone during cell plate formation. (A) Table summarizing loci putatively encoding Sec1/munc18-like (SM) family proteins and their identity to the cytokinetic Sec1 protein KEULE of Arabidopsis at the protein level. The locus identifiers for two versions of the *P. patens* genome assembly are given. For each predicted gene the number of exons and the presence of indicators of gene expression in protonemal tissue are given. (B) Comparison between the exon (black boxes) and intron (connecting chevrons) structure of the gene encoding Arabidopsis KEULE and the putative *P. patens* orthologue expressed in protonema. (C) Yeast-two-hybrid interaction assay between Sec6, PpKEULE, and the PpKEULE C1 domain (illustrated at the top). Proteins were fused with the Gal4 activation domain (AD) and binding domain (BD) and their different combinations were co-expressed in yeast strains that were tested for growth on reporter media lacking histidine in combination with increasing amounts of the competitive inhibitor 3-AT. (D) Sec6 and PpKEULE co-visualized throughout cell division in a cell expressing Sec6-mCherry and PpKEULE-GFP. The arrowheads indicate Sec6-labelled regions at the expanding edge of the phragmoplast showing less associated KEULE-GFP compared to the central regions. An enlarged view of the bracketed area is depicted in the inset where the arrow points to Sec6 located at the cortex without associated PpKEULE-GFP signal. Images are maximum z-projections of 3 confocal planes spaced 0.5 μm apart. Scalebar, 5 μm.

Introns in the PpKEULE gene were slightly longer, however this is in line with average intron length being bigger in moss compared to Arabidopsis (Rensing et al., 2005). The concurrent intron-exon structure of the genes of both species signifies that they are derived from a gene present in the last common ancestor of mosses and higher plants.

In Arabidopsis, the interaction domain of KEULE with Sec6 was narrowed to a C-terminal portion of the protein designated C1 (Wu et al., 2013). The high degree of sequence conservation allowed us to delineate the same domain in PpKEULE and test it for its ability to interact with Sec6. In a reciprocal yeast 2-hybrid assay with *P. patens* Sec6, KEULE and the KEULE-C1 domain, we found a strong interaction between Sec6 and both PpKEULE and PpKEULE-C1 when Sec6 was fused to the activation domain of the Gal4 transcription factor. When Sec6 was fused to the DNA-binding domain of Gal4 only a weak interaction with the PpKEULE-C1 domain was found (Figure 5C). Possibly, in this case, the binding partners are in a less favourable configuration for reconstituting Gal4 function since a similar result was reported for the interaction of Arabidopsis proteins (Wu et al., 2013). Because PpKEULE interacts with Sec6 in a fashion indistinguishable from its orthologue in Arabidopsis, the two proteins could form a conserved module regulating cell plate membrane fusion. To investigate where and when in *P. patens* Sec6 and PpKEULE may interact, we visualized Sec6-mCherry together with endogenous PpKEULE tagged with GFP (Figure S3). We chose to fuse GFP to the C-terminal end, since this yielded a functional fusion protein for Arabidopsis KEULE (Steiner et al., 2016). While absent from Sec6-labelled regions during prophase and metaphase, PpKEULE-GFP became rapidly enriched at the site of Sec6-mCherry at the onset of cytokinesis around the time that localized membrane deposition is first observed at overlaps (Figure 5D; Movie S5). During radial expansion of the phragmoplast, appearing sites of Sec6 localization at the leading zone, likely corresponding with newly formed overlaps with yet little accumulated vesicles (de Keijzer et al., 2017), had low PpKEULE-GFP signal associated. (Figure 5D, arrowheads). At the cortical zone marked by the labelled exocyst subunit no corresponding PpKEULE signal was found (Figure 5D, arrow). Taken together, the co-localization pattern suggested that physical association between Sec6 and PpKEULE in a cytokinetic context is both spatially and temporally regulated.

To investigate the functional importance of Sec6 for correct KEULE localization and cell plate assembly in general we sought ways to reduce or abolish cellular Sec6 levels. We failed to isolate a Sec6 knock-out mutant, suggesting that Sec6 is an essential protein for cell proliferation, as has been reported by van Gisbergen et al., (2018). We therefore aimed to generate an inducible knock down of Sec6 transcript levels using inducible RNAi in cells expressing either vesicle marker SCAMP4-mCherry or PpKEULE-GFP. Induction of Sec6 RNAi caused a large reduction of moss colony expansion and caulonemal cells were absent (Figure S4A and S4B). Nonetheless, chloronemal cells with induced RNAi showed tip growth albeit at a decreased rate and with a more bulbous tip shape (Figure S4C).

This may suggest that some functional exocyst complexes are present, or that a minimal amount of tip growth can occur in the absence of Sec6. Three days after induction of RNAi knockdown, finalized cell walls separating two daughter cells showed various morphological defects (Figure S4D). To understand during which stage of cell plate assembly these defects could arise, we quantified cell plate expansion, membrane deposition, and KEULE localization. Cell plate expansion continued at near wild type levels but levels of SCAMP4-mCherry and PpKeule-GFP at the phragmoplast midzone were decreased after RNAi induction (Figure 6; Movie S6 and S7). Later in cytokinesis, protein levels were similar to non-induced cells. Our RNAi experiments thus suggest that reduction of Sec6 functioning at the midzone reduces membrane build up during early cytokinesis, a time at which Sec6 levels peak in unperturbed cells (Figure 1A).

**Figure 6.**
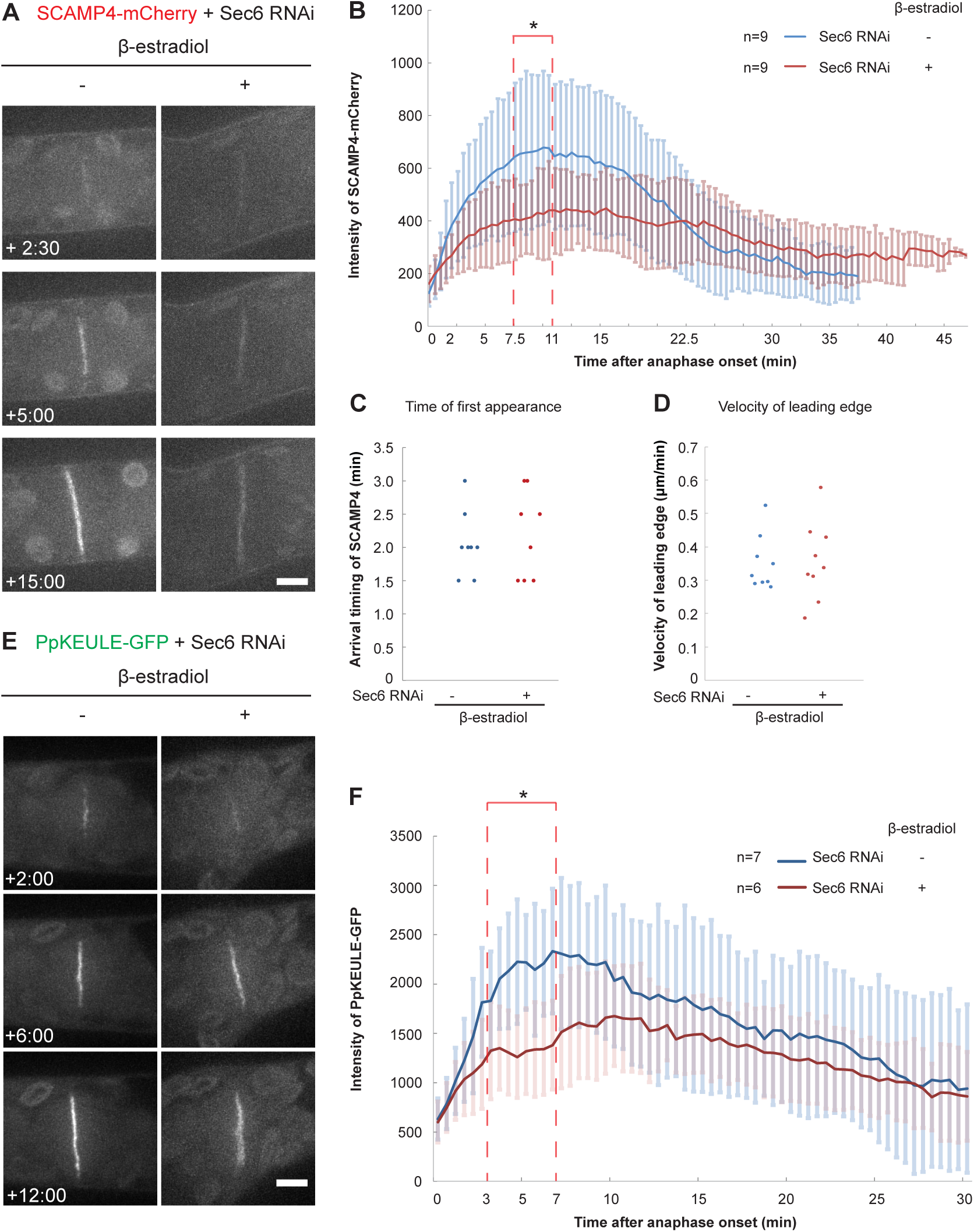
Initial recruitment of membrane materials and PpKEULE is reduced upon Sec6 silencing. (A) Snapshots of dividing cell with membrane marker SCAMP4-mCherry in Sec6 RNAi without or with RNAi induction by β–estradiol treatment. The intensity of SCAMP4-mCherry is depicted with the same contrast settings. Time with respect to anaphase onset (t=0) is indicated in min:sec. Images are maximum z-projections of 3 planes spaced 0.5 μm apart acquired in the central plane. Scale bar =5 μm. (B) Average intensity of SCAMP4-mCherry quantified throughout cytokinesis. The error bars represent SD of each time point. * indicates the significant difference (P <0.05) between control and Sec6 RNAi from 7.5 to 11 min in Student’s two-tailed t-tests. (C) The onset of membrane arrival in Sec6 RNAi background with or without induction. Anaphase onset is set up as t=0. n=8 for both treatments. (D) The expansion rate of the cell plate in Sec6 RNAi background with or without induction. n=8 for both control and Sec6 RNAi. (E) The Localization of PpKEULE-GFP in Sec6 RNAi background. Representative snapshots are shown in the early, middle, and late phase of cytokinesis. Images are maximum z-projections of 3 planes spaced 0.5 μm apart acquired in the central plane. KEULE-GFP is depicted using identical contrast settings among RNAi and control cells. Scale bar =5 μm. (F) Average PpKEULE-GFP intensity throughout cytokinesis. The error bars represent SD of each time point. * indicates the significant difference (P <0.05) between control and Sec6 RNAi from 3 to 7 min in Student’s two-tailed t-tests.

## Discussion

Our cytokinesis-oriented localization screen of exocyst subunits in moss revealed two localization patterns that have not been previously described in plants. Firstly, exocyst subunits localized to microtubule overlaps from early mitosis onwards, and secondly, they formed a cortical ring prior to cell plate attachment. Interestingly, the two localization patterns overlapped in time yet differed in subunit composition. On overlaps, Sec6 arrival preceded Sec5 and Sec3 whilst this was not evident for the cortical localization. The systematic design of the conducted screen thus provides some of the clearest evidence for sequential assembly of the exocyst complex to date. So although biochemical purification and live cell data show that eight exocyst subunits in yeast form a stable complex, our work shows that subunits can have individual localization patterns as well (Hála et al., 2008; Heider et al., 2016; Picco et al., 2017).

In most plant cells a cortical ring of microtubules, the preprophase band (PPB), is thought to demarcate a functionalized section of the cortex, called the cortical division zone (CDZ), to which cell plate expansion is directed during cytokinesis (van Damme, 2009; Müller et al., 2009; Stöckle et al., 2016). Known components of the CDZ do however assemble into ring-like structures in mutants that are unable to form a PPB (Schaeffer and Bouchez, 2010). Other cells, including moss protonemal cells, lack a PPB altogether but cytokinesis does involve the formation of a cortical band with established CDZ components, including myosins and kinesins, prior to cell plate attachment (Schmiedel et al., 1981; Doonan et al., 1985; Otegui and Staehelin, 2000; Hiwatashi et al., 2008; Wu et al., 2011; Nakaoka et al., 2012; Miki et al., 2014; Wu and Bezanilla, 2014; Lipka et al., 2014; Schaefer et al. 2017; Kosetsu et al., 2017;,). Given the early cortical localization of the exocyst observed in moss, it will be interesting to find out if the exocyst and vesicle trafficking have a role in the establishment or maintenance of the CDZ in absence of a PPB, or whether its function is limited to attachment of the cell plate. Analogies may be present with fission yeast cytokinesis where exocyst localization and vesicle fusion to the division site appear to precede cleavage furrow ingression (Wang et al., 2016). It is surprising that no exocyst subunits have been reported as components of the CDZ in higher plants (Vukašinović and Žárský, 2016; Borucand van Damme, 2015). Characterization of exocyst localization in plants is however impeded by the large number of subunits homologues, particularly of the Exo70 subunit.

Local accumulations of exocyst subunits were first described for their role in polarized exocytosis at the plasma membrane (TerBush and Novick 1995; Hazuka et al., 1999). However, exocyst subunits also localize to internal membrane compartments to assist in autophagosome formation (Kulich et al., 2013; Bodemann et al., 2011). Moreover, in budding and fission yeast it was demonstrated that the deliberate targeting of exocyst subunits to mitochondria reroutes cellular trafficking towards these organelles (Luo et al.,, 2014; Wang et al., 2016). These observations lead us to propose that a local pool of exocyst subunits on microtubule overlaps in moss may stimulate the immobilization and fusion of vesicles to initiate cell plate assembly. However, in all mentioned examples local pools of exocyst subunits are bound to membranes whereas Sec6 in our studies localized to non-membranous microtubule overlaps. It is therefore not self-evident how Sec6 on overlaps may initiate membrane-membrane interactions for homotypic vesicle fusion even when it seeds the assembly of a complete exocyst complex. Possibly, vesicles arrive at the phragmoplast midzone in a fusion-incompetent state and formation of enabled SNARE complexes involves interactions between SNAREs, KEULE and exocyst subunits. In fact, Sec6 appears to be at the centre of an interaction network since SEC6-homologues in diverse systems interact with both SNARE- and SM proteins apart from other exocyst subunits (Hong and Lev, 2014; Hashizume et al., 2009; Morgera et al., 2012; Dubuke et al., 2015; Siviram et al., 2005, 2006). Cytosolic levels of Sec6 may be insufficient to establish these regulatory molecular interactions on vesicles whilst concentrated Sec6 on overlaps could generate a local pool of fusion-competent vesicles near overlaps. Our finding thus suggests how vesicle fusion activity may be regulated in space, an aspect that so far remained unresolved. Interestingly, *Pp*KEULE does not localize to cortical exocyst subunits prior to cell plate attachment. It may thus be exclusively required to regulate vesicle fusion activity in the phragmoplast midzone during cytokinesis.

Very unique about Sec6 during moss cytokinesis is its early localization to microtubule overlaps in complete absence of membranes. Other exocyst subunits may be involved in binding to overlaps but not Sec5 and Sec3 which arrive later, around the time of membrane accumulation. Observations in mammalian cells suggest that microtubule overlaps may have a more common role in controlling vesicular transport. Through an interaction with centralspindlin the protein MICAL3 first localizes to the spindle midzone starting in anaphase and later targets Rab8a positive vesicles to the midbody (Liu et al., 2016). Centralspindlin does however transfer from microtubule overlaps to an adjacent ring-shaped bulge where also exocyst subunits are localized (Hu et al., 2012; Gromley et al., 2005). It is therefore not clear how intimate the association between overlaps and vesicles is in these cells (Green et al., 2013). Moss does not have centralspindlin but other midzone associated proteins like MAP65 may have a function in Sec6 recruitment to microtubule overlaps. Such physical interactions between cytoskeletal regulators and exocyst subunits were reported in various other systems (van Gisbergen et al., 2018; Gromley et al., 2005; Zuo et al., 2006; Oda et al., 2015). Although MAP65 silencing and Δ*kin4-Ic* experiments clearly showed that the length of microtubule overlaps template Sec6 localization, the protein was also present away from overlaps and localized to the full cell plate at later stages of cytokinesis. We hypothesize that Sec6 and possibly other proteins on overlaps are gradually incorporated in a structure that also drives membrane accumulation away from overlaps. This structure may represent the Cell Plate Assembly Matrix (CPAM) that is visible in EM micrographs and from which microtubule overlaps are excluded (Seguí-Simarro et al., 2004). Cell plate dimensioning and assembly defects in cells lacking Kin4 do however show that overlap length is important for templating cell plate initials, likely in part through patterning of the exocyst complex (de Keijzer et al., 2017).

Our findings provide the first indication that molecules driving membrane fusion processes are at least in part associated with microtubule overlaps during plant cytokinesis. The formation of intracellular septa in general is a complex task that requires spatial control over vesicle fusion to make a *de novo* plasma membrane. The involvement of multiple tethering complexes has been reported in various eukaryotes (see Neto and Gould, 2011; Wang et al., 2016; Rybeck et al., 2014). In part these complexes may act redundantly, which hinders their functional characterization. It will be interesting to learn how positioning tethering complexes may fine tune cell plate formation on the ultrastructural level. Differential localization of TRAPPII and exocyst complexes was reported during fission yeast cytokinesis and the two complexes in Arabidopsis where present sequentially at the developing cell plate (Wang et al., 2016; Rybak et al., 2014). The molecular and light microscopic tools available in the moss, together with the identification of overlaps as sites of vesicle accumulation, will help to resolve how multiple tethering complexes together shape the cell plate.

## Materials and methods

### Plasmids and cloning procedures

All plasmids used throughout this study are listed in Supplemental table 2. For construction of GFP/mCherry tagging constructs for Sec6, Sec3a, Sec3b, Sec3c, Sec5a, Sec5b, Sec5d, and PpKEULE, regions of approximately 1 kb before and after the stop codon of the encoding gene were amplified by PCR using primers listed in Supplemental table 1. The PCR fragments were digested with restriction enzymes indicated in supplemental table 1and ligated into correspondingly digested peGFP-NPTII or pmCherry-LoxP-BsdR vector (de Keijzer et al., 2017).

For β-estradiol inducible RNAi, targeting Sec6 transcript, we used the system published by Nakaoka et al. (2012). A ~500 bp fragment of the coding sequence of Sec6 (figure S5) was amplified by PCR from a cDNA library derived from protonemal tissue and cloned into gateway entry plasmid pENTR-D-TOPO. The fragment was subsequently introduced into silencing vector pGG626 via a Gateway LR reaction.

To generate the expression constructs used for yeast-two-hybrid assays, first the coding sequences of Sec6, PpKEULE, and the PpKEULE-C1 domain were PCR-amplified from a *P. patens* cDNA library derived from protonemal tissue. The PCR products were then introduced into the pENTR-D-TOPO vector and subsequently subcloned into destination plasmids pDEST22 and pDEST32 (Invitrogen) via Gateway LR reactions.

### *P. patens* growth conditions and transformation

*P. patens* tissues were routinely grown on BCDAT plates under continuous light. Plasmids were linearized and introduced into the *P. patens* genome by homologous recombination using PEG-mediated protoplast transformation (Nishiyama et al., 2000). Correct insertion events were characterised by PCR (Figure S3). Knock-down in RNAi lines upon induction was verified by quantitative RT-PCR (Figure S5). Characteristics of generated moss lines and their use throughout the study are summarized in Supplemental table 3. For imaging, protonemal tissue was grown for 5 to 7 days on BCD medium in glass bottom dishes (Yamada et al., 2016).

### Identification of *P. patens* Sec1 homologues

Proteins of the Sec1/Munc18 family were identified by BLAST using the Arabidopsis KEULE protein sequence as input against the predicted proteins of *P. patens* genome assemblies version 1.6 and 3.1 and by keyword search in the Phytozome 10.3 browser (www.phytozome.org). Expression in protonema was assessed by verifying the presence of ESTs derived from protonemal tissue and using the *Physcomitrella* eFP browser (Ortiz-Ramírez et al., 2016).

### Yeast two-hybrid assay

Yeast 2-hybrid assays were performed with a split Gal4 transcription factor system using the His3 gene as reporter (James et al., 1996). For this, pDEST22/32-based constructs (supplemental table 2) were transformed into yeast strain PJ69-4a or PJ69-4α by PEG-mediated transformation. Positive transformants with minimal background reporter activity were selected on double dropout medium (–Leu–His or –Trp–His) with different concentrations of 3-amino-1,2,4-triazole (3-AT) present to increase histidine-dependent growth stringency. Selected clones were then allowed to mate and resulting diploids were selected on –Leu–Trp plates. With surviving the cells a yeast 2-hybrid assay was then performed on triple dropout medium (–Leu–Trp–His) with increasing amounts of 3-AT present.

### Fluorescence microscopy and staining

All live cell imaging was performed on a Roper spinning disk microscope system composed of a Nikon Ti eclipse body, Yokogawa CSU-X1 spinning disc head and Photometrics Evolve 512 camera. All imaging was conducted with a 100x Plan Apo VC oil immersion objective (NA 1.40), using a 1.2x post-magnification fitted before the camera. The GFP and citrine probes were exited using 491 nm light generated by a Cobolt Calypso50 laser and their emitted light was bandpass filtered at 497-557 nm. For FM4-64 and mCherry 561 nm excitation light generated by a Cobolt Jive50 laser was used in combination with bandpass filtering at 570-620 nm. During image digitization a camera electron multiplication gain of 300 was employed and typical exposures were 800-1000 ms for GFP, citrine and FM4-64 probes and 1000-2000 ms for the mCherry probe. FM4-64 was dissolved in dH2O at a final concentration of 10 μM and added to cells at the moment of nuclear envelope breakdown as described (Kosetsu et al., 2013).

### Image analysis

FIJI (Schindelin et al., 2012) was used for all image analysis and processing. Figures were prepared in Adobe Illustrator CC 2015.

### Analysis of Sec6 RNAi mutant phenotype

To quantify the accumulation of cell plate membrane material and PpKEULE upon Sec6 RNAi silencing, we imaged SCAMP4-mCherry and PpKEULE-GFP throughout cytokinesis at a 30 second intervals. Images were taken in the central plane of the dividing cell at 3 confocal planes spaced 0.5 μm apart. Maximum projections made along the z-axis were then used to measure the average intensity of SCAMP4-mCherry and KEULE-GFP using a 5 pixel wide line manually drawn along the division plane for each time point. Simultaneously, the mean level of cytosolic background fluorescence was recorded in an area right next to the cell division site. For each time point, the background fluorescence level was subtracted from the obtained level of fluorescence at the division plane. The expansion rate of the cell plate (Figure 6D) was calculated as described by de Keijzer et al. (2017).

## Acknowledgements

We would like to thank Henk Kieft for technical assistance.

## Data availability

For gene annotation, Cosmoss (www.cosmoss.org/) and the Phytozome 10.3 browser (www.phytozome.org) were used. Gene expression data were obtained from the *Physcomitrella* eFP browser (bar.utoronto.ca/efp_physcomitrella/cgi-bin/efpWeb.cgi).

